# Biofilm Formation by *Staphylococcus aureus* is Triggered by a Drop in the Levels of the Second Messenger cyclic-di-AMP

**DOI:** 10.1101/2020.01.31.929125

**Authors:** Adnan K. Syed, Christopher R. Vickery, Taliesin Lenhart, Eliza Llewellyn, Suzanne Walker, Richard Losick

## Abstract

The bacterial pathogen *Staphylococcus aureus* forms multicellular communities known as biofilms in which cells are held together by an extracellular matrix. The matrix consists of repurposed cytoplasmic proteins and extracellular DNA. These communities assemble during growth on medium containing glucose, but the intracellular signal for biofilm formation was unknown. Here we present evidence that biofilm formation is triggered by a drop in the levels of the second messenger cyclic-di-AMP. Previous work identified genes needed for the release of extracellular DNA, including genes for the cyclic-di-AMP phosphodiesterase GdpP, the transcriptional regulator XdrA, and the purine salvage enzyme Apt. Using a cyclic-di-AMP riboswitch biosensor and mass spectrometry, we show that the levels of the second messenger drop during biofilm formation in a glucose-dependent manner and that the drop is prevented in mutants of all three genes. Importantly, we also show that expression of the “accessory gene regulator” operon *agr* is under the positive control of cyclic-di-AMP and that an *agr* mutation, which is known to promote biofilm formation, bypasses the block in biofilm formation and eDNA release caused by a *gdpP* mutation. We conclude that the effect of the glucose-dependent drop in c-di-AMP levels is principally mediated by a reduction in *agr* expression, which in turn promotes biofilm formation.

## Introduction

Biofilms are microbial communities in which cells adhere to each other and to a surface by an extracellular matrix composed of DNA, proteins, and/or polysaccharides. These matrix components are generally produced by the microbes in the community either through dedicated pathways, as in the cases of curli and cellulose production in *Escherichia coli* and TasA and the EPS exopolysaccharide in *Bacillus subtilis*, or through lysis of a subset of cells and recycling of cytoplasmic components, as in the cases of *Pseudomonas aeruginosa* and *Staphylococcus aureus* (Chapman et al. 2002; Serra et al. 2013; Branda et al. 2006; 2005; Whitchurch et al. 2002; Turnbull et al. 2016; Gloag et al. 2013; Mann et al. 2009; Foulston et al. 2014; Dengler et al. 2015). The biofilm state confers resistance to a variety of assaults including from predators, competitors, and the immune system (DePas et al. 2014; Stewart and Costerton 2001; Høiby et al. 2010; Bhattacharya et al. 2018; Hanke and Kielian 2012). Biofilm formation often also plays an essential role in recalcitrant infections caused by bacteria such as *S. aureus* by contributing to antibiotic tolerance and evasion of immune cells (Bjarnsholt 2013; Römling et al. 2014).

*S. aureus* strains, and methicillin-resistant strains (MRSA) in particular, form biofilms by recycling cytoplasmic proteins and chromosomal DNA (Mann et al. 2009; Foulston et al. 2014; McCarthy et al. 2015). The proteins, which are positively charged at the low pH under which biofilm formation takes place (e.g. fermentation during growth on medium containing glucose) associate with the negatively charged cell surface (Foulston et al. 2014). Evidence indicates that the recycled chromosomal DNA (referred to as extracellular DNA or eDNA), forms an electrostatic net with the protein coated cells holding them together (Dengler et al. 2015; Kavanaugh et al. 2019). Biofilm formation by methicillin-sensitive strains of *S. aureus* (MSSA) generally rely on a secreted exopolysaccharide known as Polysaccharide Intercellular Adhesin (PIA), which is produced by the *ica* operon (Jefferson et al. 2003).

Using a comprehensive and unbiased genome-wide transposon sequencing approach, we previously identified several genes that are required for the release of eDNA from *S. aureus* cells during glucose-dependent biofilm formation (DeFrancesco et al. 2017). Among these were *gdpP*, which encodes the cyclic-di-AMP (c-di-AMP) phosophodiesterase GdpP, *xdrA*, which specifies the transcription regulatory protein XdrA, and *apt*, the gene for the adenine phosphoribosyltransferase Apt. Mutants of all three genes exhibited marked defects in the release of eDNA and biofilm formation. Analysis using HPLC showed that the levels of c-di-AMP decreased during growth of the wild type on biofilm-inducing medium [Tryptone Soy Broth (TSB) containing glucose] whereas the levels rose in cells mutant for GdpP. These findings raised the possibility that biofilm formation in *S. aureus* is triggered by a drop in c-di-AMP levels.

Here, we present evidence in support of this hypothesis. First, we demonstrate that dependence on eDNA is a shared feature of biofilm formation by multiple strains of *S. aureus*, including an MSSA strain in which biofilm formation is dependent on PIA. Second, the dependence of eDNA release on GdpP is shared among biofilm-forming strains. Third, using a c-di-AMP sensitive riboswitch biosensor and mass spectrometry, we confirm that the levels of c-di-AMP drop under biofilm-inducing conditions, and we additionally show that the drop is prevented in null mutants of *xdrA* and *apt* as well as in a *gdpP* mutant. Fourth, we report that overexpression of *gdpP* reverses the block in eDNA release and in biofilm formation caused by the *xdrA* mutation. Finally, and importantly, we show that expression of the “accessory gene regulator” operon *agr* and genes under its control is dependent on c-di-AMP and that the decrease in *agr* expression is principally responsible for triggering eDNA release and biofilm formation in response to the drop in second messenger levels (Recsei et al. 1986).

## Results

### eDNA release is a common feature of biofilm formation in diverse *S. aureus* strains

Because biofilm formation is studied in different laboratories using a variety of strains, we began by asking whether eDNA release is a common feature of biofilm formation by *S. aureus*. Accordingly, we investigated biofilm formation using: HG003, the standard strain used in the investigators’ laboratories, SH1000, RN1, MN8, LAC, and Newman (Herbert et al. 2010; O’Neill 2010; Voyich et al. 2005; Duthie and Lorenz 1952; Maira-Litrán et al. 2002). In our hands, four of these strains (HG003, SH1000, RN1, and MN8) formed robust biofilms as judged by biomass and did so in a glucose-dependent manner (Figure 1). Moreover, as judged by light microscopy HG003, SH1000, RN1, and MN8 formed large clumps of cells that adhered to each other (Figure 2). Finally, all four strains were robust in eDNA release during biofilm formation, and the integrity of the resulting biofilms was dependent on the eDNA in that the biofilms (Figure 1) and cell clumps (Figure 2) were substantially dispersed by treatment with the enzyme DNase I. In contrast, the LAC and Newman strains were relatively unresponsive to growth on glucose as judged by all three criteria, namely, eDNA release, biofilm formation, and cell clumping (Figures 1 and 2). We conclude that eDNA release and dependence on eDNA for biofilm formation and integrity are characteristic of strains that form biofilms in a glucose-dependent manner.

**Figure 1.**
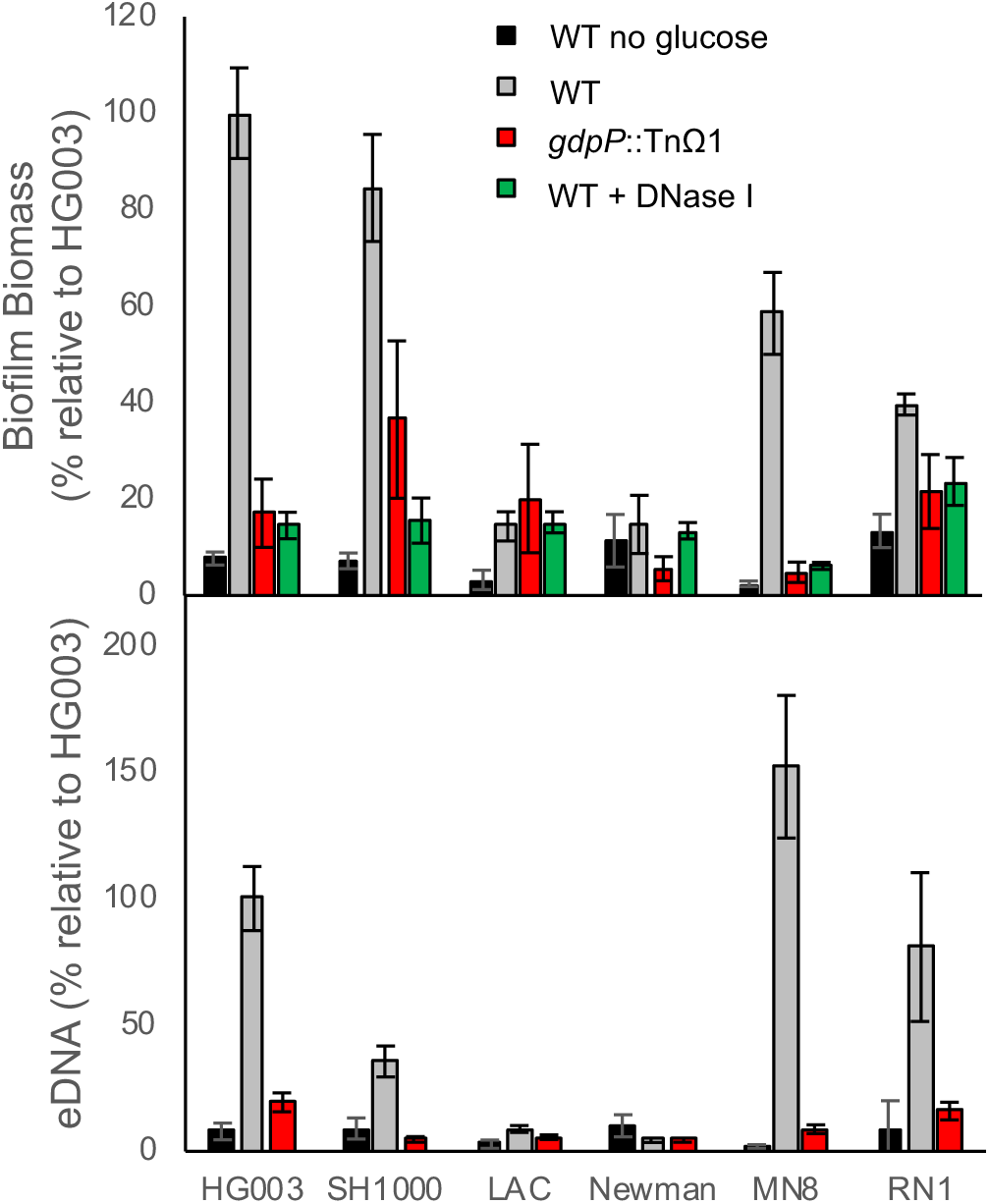
eDNA release and dependence on *gdpP* is a shared feature of biofilm formation by multiple strains of *S. aureus*. The indicated strains were tested for their ability to form glucose- and DNA- dependent biofilms (top) and to release eDNA (bottom). *gdpP*::TnΩ1 (red) was transduced into each strain and the mutant strains were tested for their ability to form glucose-dependent (top) biofilms and release eDNA (bottom). DNase I was added to established biofilms for 1 hour prior to harvesting to test the dependence on DNA (green).

**Figure 2.**
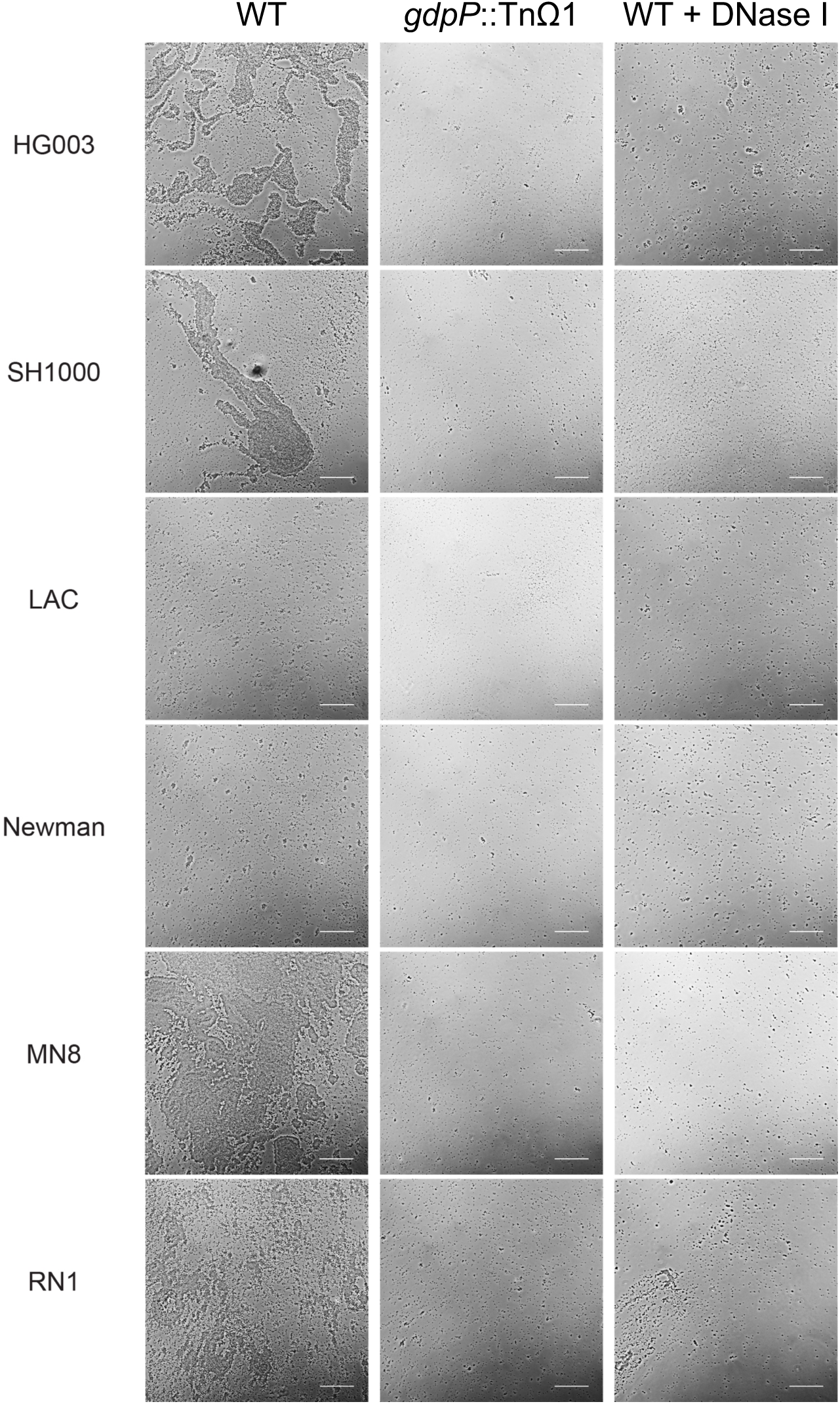
Cell-cell adhesion is dependent on eDNA and blocked by *gdpP*::TnΩ1. Microscopy was performed on gently resuspended biofilms. Scale bars are 200 μm in length.

Of special interest were the results with strain MN8 as this strain is known to produce biofilms in a manner that depends on the exopolysaccharide PIA. Still, in our hands, MN8 was found to release eDNA during biofilm formation and the integrity of the resulting biofilms was strongly dependent on eDNA (Figures 1 and 2).

### Dependence on the c-di-AMP phosphodiesterase GdpP is a general feature of eDNA release during biofilm formation

Our previous work has shown that in strain HG003, eDNA release during biofilm formation and biofilm formation itself are significantly impaired by mutation of the gene for the c-di-AMP phosphodiesterase *gdpP* (DeFrancesco et al. 2017). To investigate whether this is generally the case, we transduced the transposon insertion *gdpP*::TnΩ1 from HG003 into SH1000, RN1, MN8, LAC, and Newman. The four strains that formed robust biofilms in a glucose-dependent manner (namely, HG003, SH1000, RN1, and MN8) all exhibited marked decreases in eDNA release, biofilm formation, and cell clumping when the *gdpP* mutation was introduced (Figures 1 and 2). As expected, the *gdpP* mutation had little or no effect on the biofilm-impaired strains LAC and Newman (Figures 1 and 2). These data are consistent with our hypothesis that *gdpP* is generally needed for the release of eDNA during glucose-dependent biofilm formation.

### c-di-AMP levels drop during biofilm formation

Previously, using HPLC, we found that c-di-AMP levels were lower in biofilm cells than in cells grown in the absence of glucose, leading to the hypothesis that eDNA release and biofilm formation are triggered by a drop in the levels of the second messenger (DeFrancesco et al. 2017). To facilitate further tests of this hypothesis, we turned to the use of a riboswitch reporter for c-di-AMP created, and kindly provided to us, by Ming Hammond (Kellenberger et al. 2015). The riboswitch is attached to a Spinach2 aptamer that binds to a dye that fluoresces when the riboswitch is bound by c-di-AMP, allowing a fluorescence readout of relative changes in c-di-AMP levels in living cells over time. In keeping with our previous observations, we observed a glucose-dependent drop in c-di-AMP levels commencing at 6 to 8 hours after inoculation and persisting for the duration of the experiment (Figure 3A). In contrast, cells grown in TSB medium without added glucose showed little to no change in the level of c-di-AMP during the course of the experiment (Figure 3A). HPLC-mass spectrometry was used to verify that the level of c-di-AMP did decrease in response to growth on glucose (Table S2). Furthermore, we found no significant change in the transcript levels of the diadenylate cyclase *dacA* or c-di-AMP phosphodiesterase *gdpP* genes (relative to those for the house keeping gene *hu*, as an internal control) in response to glucose (Table S3).

**Figure 3.**
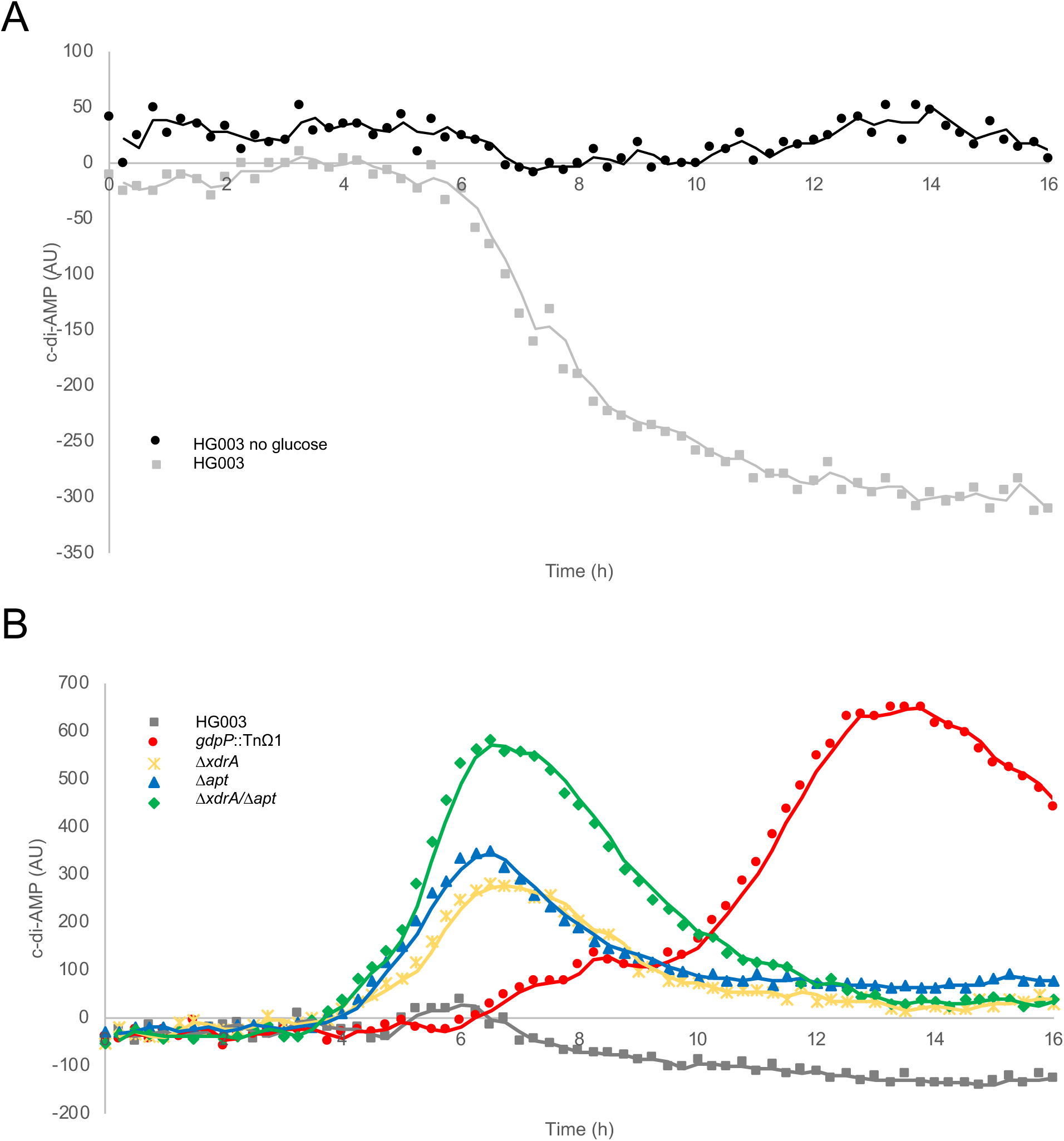
Monitoring cyclic-di-AMP levels during biofilm formation using a riboswitch biosensor. Cells containing a cyclic-di-AMP sensitive riboswitch biosensor were grown under conditions that do not permit biofilm formation (no added glucose, black) or biofilm promoting conditions (plus glucose, grey). (A) cyclic-di-AMP levels dropped between 6 and 8 hours in a glucose-dependent manner. (B) *gdpP*::TnΩ1 (red), Δ*xdrA* (yellow), Δ*apt* (blue), Δ*xdrA*/Δ*apt* (green) mutations caused elevated levels of c-di-AMP compared to that of the wild-type HG003 (grey) under biofilm-forming conditions. The baseline has been normalized to HG003 grown with no added glucose.

### The drop in c-di-AMP levels is also dependent on two other genes needed for eDNA release and biofilm formation

The *gdpP* gene was identified in a previous unbiased transposon screen for insertion mutants impaired in eDNA release during biofilm formation (DeFrancesco et al. 2017). The transposon screen revealed two additional genes in which mutations markedly impaired eDNA release and biofilm formation. These were *xdrA*, which encodes a DNA-binding protein, and *apt*, which codes for an adenine phosphoribosyltransferase involved in a purine salvage pathway. We wondered whether *xdrA* and *apt* were also needed for the drop in c-di-AMP levels and whether this could be the basis, in whole or in part, for their involvement in biofilm formation. Accordingly, we used null mutations of each gene (*ΔxdrA* and *Δapt*) and the riboswitch to investigate whether either or both genes were needed for the drop in c-di-AMP levels during biofilm formation. The results of Figure 3B show that both mutations completely prevented the drop in c-di-AMP levels seen in the wild type. Indeed, the levels of the second messenger rose somewhat in both mutants, although not to the extent observed for the *gdpP* mutant. That the *ΔxdrA* and *Δapt* mutations blocked the drop in c-di-AMP levels was independently confirmed by HPLC-mass spectrometry (Table S2).

Next, we asked whether the effects of the mutations could be reversed by overexpression of the gene for *gdpP* using a plasmid harboring the phosphodiesterase gene (p*gdpP*). The results were particularly striking for *ΔxdrA*. The p*gdpP* plasmid increased eDNA release approximately 5 fold for the *ΔxdrA* mutant (while also increasing eDNA levels for the wild type and the *gdpP* insertion mutant) (Figure 4A). The plasmid also markedly increased the amount of biofilm mass of the *ΔxdrA* mutant and in a manner that was sensitive to DNase I (Figure 4B). As a control, we found that a mutant version of the plasmid producing GdpP with reduced phosphodiesterase activity due to a D418A substitution (Rao et al. 2010; Griffiths and O’Neill 2012) was not able to fully restore eDNA release or biofilm formation to a *gdpP* or *ΔxdrA* mutant (Figure S1). These results are consistent with the hypothesis that eDNA release and biofilm formation are caused by a drop in c-di-AMP levels and that the *ΔxdrA* mutation exerts it effect in whole or in part by blocking the drop in second messenger levels.

**Figure 4.**
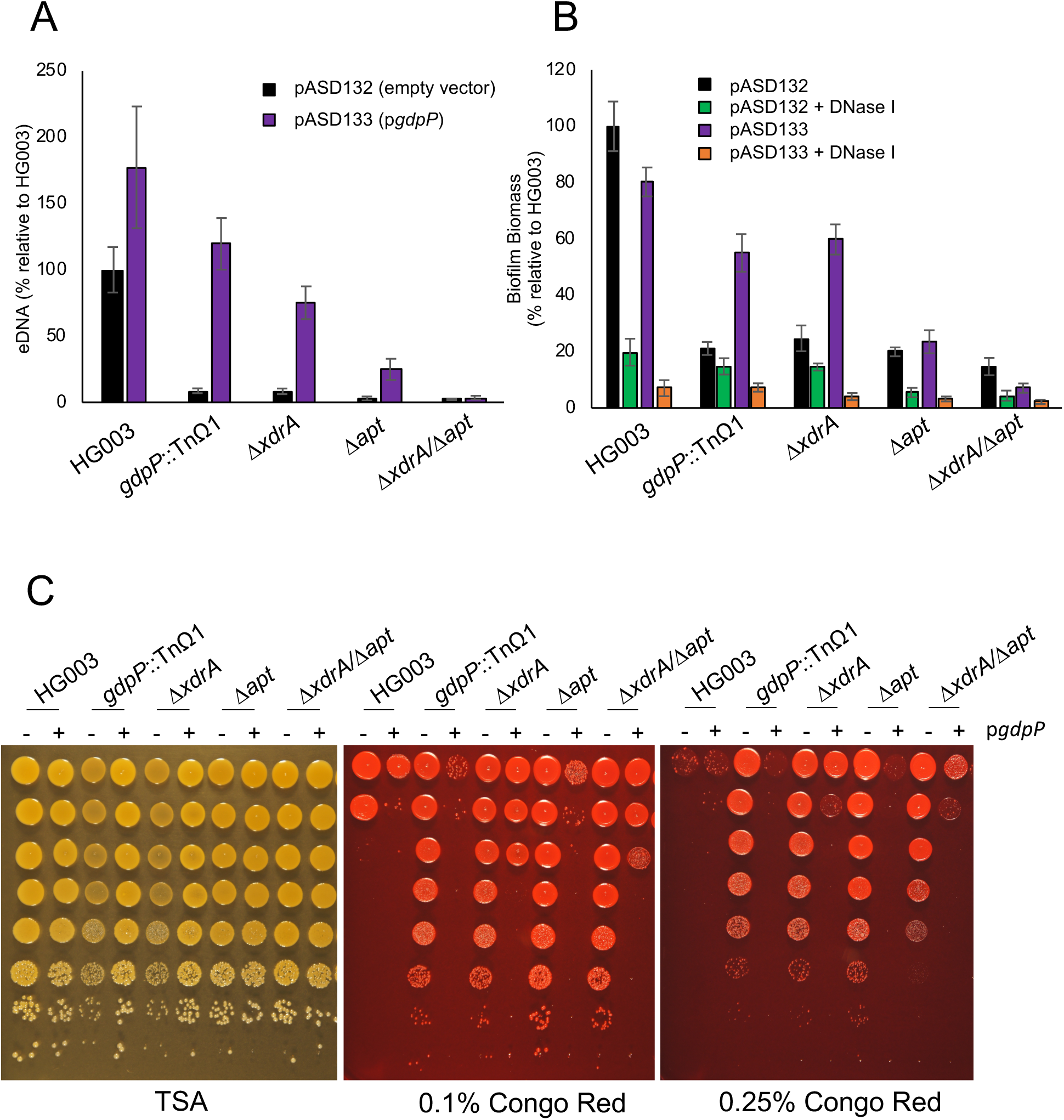
Overexpressing *gdpP* reverses the effects of an Δ*xdrA* mutation but only some of the effects of an Δ*apt* mutation. eDNA release (A) and biofilm formation of the parental strains containing an empty vector plasmid (pASD132, black) or a plasmid overexpressing *gdpP* (pASD133, purple) during glucose-dependent biofilm formation. (B) Established biofilms were tested for sensitivity to DNase I treatment (green and orange). (C) Serial dilutions of overnight cultures grown in TSB were spotted on TSA, 0.1% Congo Red, or 0.25% Congo Red and incubated overnight at 37°C prior to imaging. Strains either contained pASD132 (empty vector, -), or pASD133 (p*gdpP*, +).

The results with regard to eDNA release were also striking for the *Δapt* mutation; as seen in Figure 4A, the p*gdpP* plasmid increased eDNA levels several fold in an *Δapt* mutant. In contrast, the results with respect to biofilm formation were relatively modest; the presence of the plasmid only increased biofilm mass to a small extent. We conclude that like *xdrA* the *apt* gene contributes to eDNA release by lowering second messenger levels but that the *Δapt* mutation evidently interferes with biofilm formation in some additional manner that is not bypassed by overexpressing the c-di-AMP phosphodiesterase.

Since both *xdrA* and *apt* influence the levels of c-di-AMP, we sought to determine whether they act through the same pathway by examining the levels of the second messenger in an *ΔxdrA/Δapt* double mutant. The results show that the mutations exhibited an additive effect in raising c-di-AMP levels, suggesting that the genes modulate c-di-AMP levels through independent pathways (Figure 3B and Table S2). As seen in Figure 4, the double mutant had a severe defect in eDNA release and biofilm formation. Furthermore, we found no significant change in the transcript levels of the diadenylate cyclase *dacA* or c-di-AMP phosphodiesterase *gdpP* genes in response to mutating *gdpP*::TnΩ1, *ΔxdrA, Δapt, or ΔxdrA/Δapt*, raising the possibility that DacA or GdpP is regulated at a posttranscriptional level (Table S4).

### c-di-AMP controls *agr* expression and genes under *agr* control

The results so far raise the question of the nature of the downstream target of c-di-AMP that is responsible for triggering biofilm formation. A clue came from a previous RNA-seq analysis of the regulon controlled by the XdrA transcriptional regulator (DeFrancesco et al. 2017). As we have seen, an *xdrA* mutant has increased levels of c-di-AMP and hence genes in the regulon might be under the direct or indirect control of the second messenger. Noteworthy was that an *xdrA* mutant had elevated levels of the “accessory gene regulon” *agr* operon and its known downstream targets, RNAIII and the genes for the phenol soluble modulins (Queck et al. 2008; Koenig et al. 2004; Ji et al. 1997). Specifically, the transcripts for these genes were higher in an *xdrA* mutant than in wildtype (HG003) cells under biofilm-inducing conditions (DeFrancesco et al. 2017).

To further investigate the effect of elevated c-di-AMP levels on *agr*, we focused on *agrA*, the gene encoding the transcriptional regulator (response regulator) of the *agr* system as well as two downstream targets, RNAIII and *psmα*. We found that *agrA* transcripts were 4-5 fold elevated in both *gdpP*::TnΩ1 and in *ΔxdrA* mutants, with RNAIII and *psmα* transcript levels also being significantly elevated (relative to those for the house keeping gene *hu* as an internal control), suggesting that *agr* transcript levels are positively regulated by c-di-AMP (Tables 1, 2). To further test this hypothesis we measured transcript levels in an *Δapt* mutant and an *ΔxdrA/Δapt* double mutant (Table 1). Again, we observed elevated levels of *agrA* transcripts in both mutants. We also used p*gdpP* (pASD133) to artificially lower the amount of c-di-AMP in cells and found that in the wild type (HG003) this led to a 3.5 fold decrease in *agrA* transcript levels and a significant reduction in RNAIII and *psmα* levels (Tables 1, 2). For comparison and as pointed out above, neither biofilm promoting conditions (presence of glucose) nor mutations in *gdpP*::TnΩ1, *ΔxdrA, Δapt, or ΔxdrA/Δapt* significantly affected the transcript levels of *dacA* or *gdpP* (Tables S3, S4). These results suggest that *agrA* transcript levels and the downstream targets RNAIII and *psmα* are modulated by c-di-AMP levels.

**Table 1.**
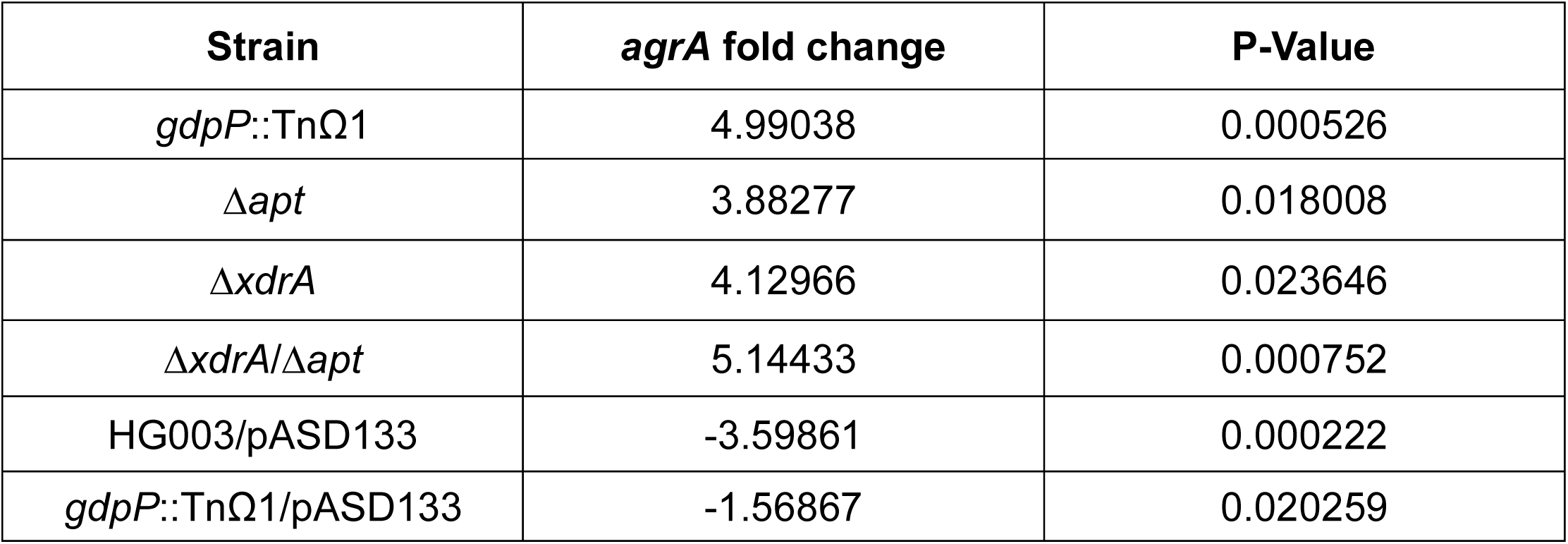
*agrA* transcript levels in the indicated strains compared to HG003

**Table 2.** RNAIII and *psmα* transcript levels in the indicated strains compared to HG003

| Strain | RNAIII |  | *psmA* |  |
| --- | --- | --- | --- | --- |
|  | Fold change | P-Value | Fold change | P-Value |
| *gdpP*::TnΩ1 | 24.98638 | 0.000142 | 5.15618 | 0.002441 |
| $\Delta xdrA$ | 17.42225 | 0.000069 | 4.13395 | 0.022991 |
| HG003/pASD133 | -5.43791 | 0.001367 | -5.33387 | 0.000678 |
| $\Delta agrA$ | -17.72638 | 0.000481 | -4.22264 | 0.000419 |
| $\Delta agrA/gdpP$::TnΩ1 | -5.68081 | 0.001672 | -5.46815 | 0.000563 |

The discovery that the *agr* system is under c-di-AMP control was pleasing as previous studies had shown that the operon is repressed during biofilm formation and that mutation of the *agr* system causes a hyperbiofilm phenotype. Indeed, as expected and in our hands, an *agrA* mutation in a HG003 background caused the release of eDNA and robust biofilm formation (Figure 5A, B). We therefore hypothesized that if elevated *agrA* transcripts in a *gdpP* mutant is preventing eDNA release and biofilm formation, then *ΔagrA* should be epistatic to *gdpP*::TnΩ1. This was indeed the case as an *ΔagrA*/*gdpP*::TnΩ1 double mutant had levels of eDNA release and biofilm formation similar to those of the wild type (HG003) (Figure 5A, B). Furthermore, using the riboswitch biosensor we found that *ΔagrA* had no effect on c-di-AMP levels (Figure 5C), in keeping with the idea that *agr* is downstream of the second messenger in the pathway governing biofilm formation. Together, these results suggest that the primary mechanism by which the drop in c-di-AMP levels under biofilm-inducing conditions triggers eDNA release and biofilm formation is by lowering the level of expression of *agr* and genes under its control.

**Figure 5.**
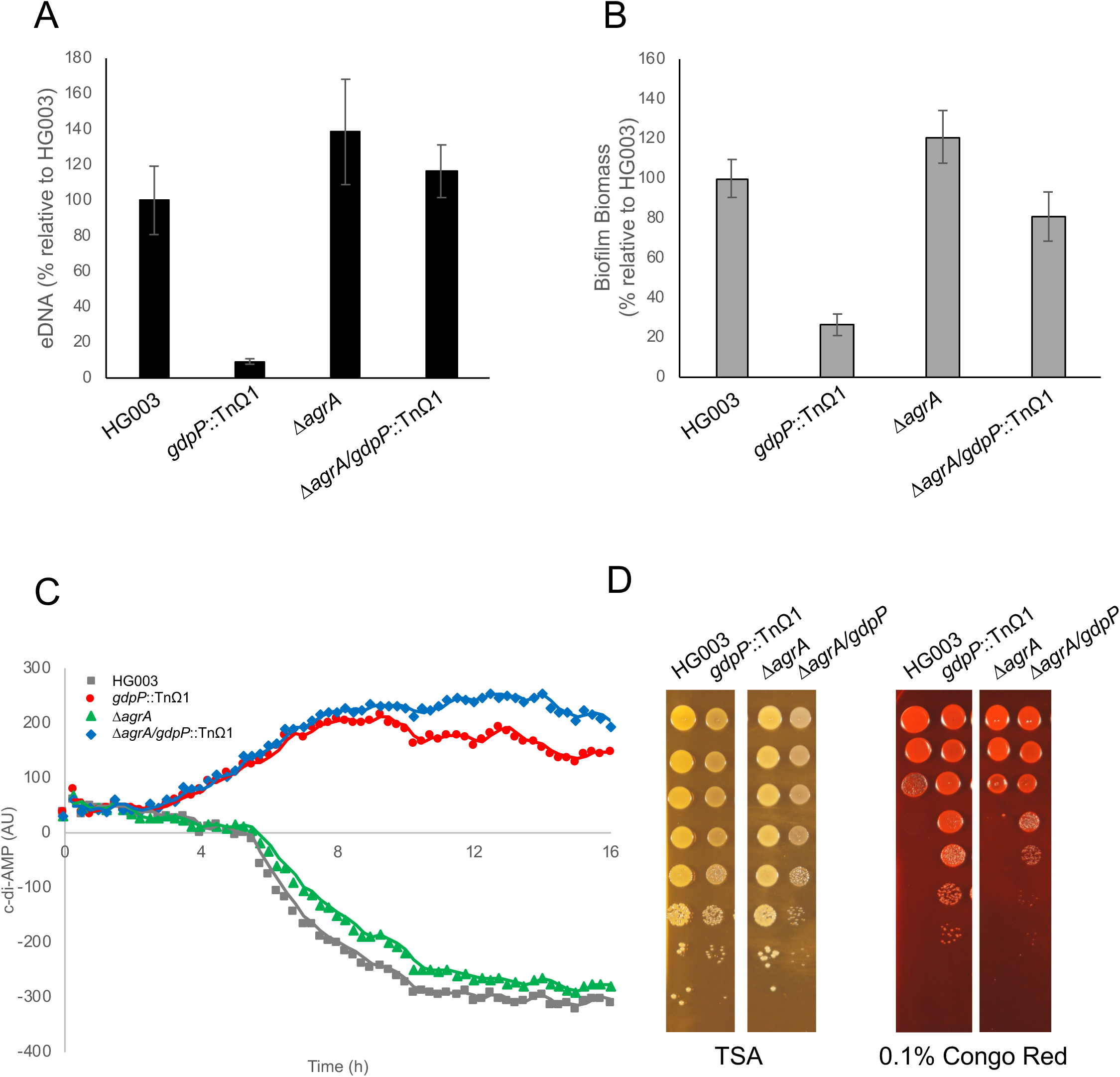
An *agr* deletion mutation is epistatic to *gdpP*::TnΩ1. eDNA release (A) and biofilm formation (B) during glucose-dependent biofilm formation. (C) Cells containing a cyclic-di-AMP sensitive riboswitch biosensor were grown under biofilm promoting conditions (plus glucose, grey) *gdpP*::TnΩ1 (red). Mutations in Δ*agrA* (green) or Δ*agrA/gdpP*::TnΩ1 (blue) do not affect the level of c-di-AMP compared to the parent strain (HG003 and *gdpP*::TnΩ1 respectively). The baseline has been normalized to HG003 no glucose (D) Serial dilutions of overnight cultures grown in TSB were spotted on TSA and 0.1% Congo Red and incubated overnight at 37°C prior to imaging.

### Overexpression of *gdpP* confers sensitivity to the dye Congo Red

As the release of eDNA during biofilm formation presumably depends on the lysis of a subset of cells, we hypothesized that mutants that are blocked in the ability to release eDNA may have sturdier cell envelopes. Previously, we have shown that wall teichoic acids protect *S. aureus* cells from lysis caused by the azo dye Congo Red, and we have also shown that mutating *gdpP, xdrA*, or *apt* all lead to increased resistance to Congo Red (Suzuki et al. 2012; DeFrancesco et al. 2017). Recently, we discovered that the target of Congo Red is LtaS, the synthetase that is responsible for the production of lipoteichoic acids (Vickery et al. 2018). We probed a HG003 transposon library with Congo Red to identify mutants with increased fitness in the presence of Congo Red and found *xdrA, gdpP*, and several other genes that were underrepresented in eDNA in the previous transposon screen (Table S5) (DeFrancesco et al. 2017). To test if *xdrA* and *apt* contribute to Congo Red resistance via their effects on c-di-AMP levels, we tested the level of resistance to the dye of null mutants of *xdrA* and *apt* containing the *gdpP* overexpression plasmid p*gdpP* (pASD133). The results in Figure 4C show that overexpressing *gdpP* restored sensitivity to Congo Red to both mutants.

To test if resistance to Congo Red was additive in an *ΔxdrA/Δapt* double mutant we determined the level of resistance at two concentrations of the dye (0.1% and 0.25%). The results show that the double mutant was no more resistant to Congo Red than either single mutant and indeed somewhat more sensitive to Congo Red than either *ΔxdrA* or *Δapt* single mutants (Figure 4C). Nonetheless, overexpression of *gdpP* restored sensitivity to Congo Red in the double mutant (Figure 4). *In toto*, these results are consistent with the idea that the Congo Red resistance of *ΔxdrA* and *Δapt* mutants is a consequence of elevated levels of c-di-AMP.

Finally, we investigated the effect of an *agrA* mutation on resistance to Congo Red. If, as we hypothesize, a *gdpP* mutation confers resistance to the dye by stimulating expression of *agr*, then an *ΔagrA* mutation should be epistatic to *gdpP*::TnΩ1. The results of Fig. 5D show that an *ΔagrA*/*gdpP*::TnΩ1 double mutant was indeed more sensitive to Congo Red than an *gdpP*::TnΩ1 single mutant although not as sensitive as an *ΔagrA* single mutant. We conclude that resistance to Congo Red as caused by elevated levels of c-di-AMP can be substantially, although not entirely, attributed to enhanced expression of *agr* (Figure 5D).

In this regard, it is interesting to note that Boyd and co-workers recently reported that cell lysis during biofilm formation can be attributed in part to decreased production of wall teichoic acids (Mashruwala et al. 2017). This fits nicely with our results with Congo Red, which as noted above is an inhibitor of lipoteichoic acid synthesis. It is known that wall teichoic acids and lipoteichoic acids contribute synergistically to cell envelope integrity (Swoboda et al. 2009). Thus, under conditions of high c-di-AMP levels and high expression of *agr*, which block biofilm formation, wall teichoic acid levels would be expected to be high, conferring resistance to Congo Red.

## Discussion

The principal contributions of this investigation are that biofilm formation and eDNA release in *S. aureus* is triggered by a drop in c-di-AMP levels and that this drop acts by lowering the expression of the *agr* operon, a negative regulator of biofilm formation. First, we have shown that eDNA is a shared feature of *S. aureus* strains that produced biofilms in a glucose-dependent manner and that in all cases eDNA release was blocked by a mutation of *gdpP*. Second, and focusing on a particular strain (HG003) that is robust in forming biofilms, we showed that the transition to a multicellular community was accompanied by a drop in c-di-AMP levels. Third, this drop was prevented by null mutations of three genes (*gdpP, xdrA* and *apt*) that were previously shown (and as confirmed here) to markedly impair eDNA release and biofilm formation. Fourth, we have shown that artificially lowering the levels of c-di-AMP by overproduction of the GdpP phosphodiesterase was suffient to reverse the phenotype of the *xdrA* mutant. Finally, we identify *agr* as the principal target for c-di-AMP and provide evidence that the effect of reduced levels of the second messenger in promoting biofilm formation and eDNA release can be largely explained by lowered expression of the *agr* operon and genes under its control.

Previous work has shown that c-di-AMP helps to maintain homeostasis during osmotic stress. Under conditions of increased uptake of osmolytes, the second messenger functions to maintain cell envelope integrity, in part though regulation of potassium transporter activity (and hence potassium uptake) (Müller et al. 2015; Commichau et al. 2018; Moscoso et al. 2015; Rocha et al. 2019; Huynh et al. 2016; Schuster et al. 2016; Devaux et al. 2018; Quintana et al. 2019; Whiteley et al. 2017; Gundlach et al. 2017; Commichau and Stülke 2018; Zeden et al. 2018; Mattingly et al. 2018; Gundlach et al. 2018). Thus, our present work reveals a new function for c-di-AMP distinct from its role in the maintainence of homeostasis.

A clue as to how c-di-AMP triggers biofilm formation has come from our discovery that a target of the second messenger is the *agr* operon. We have shown that expression of the operon and of multiple genes under its control is dependent on c-di-AMP levels. We have also shown that a null mutation of the response regulator gene in the operon, *agrA*, promotes robust biofilm formation and eDNA release even under conditions of high c-di-AMP levels (e.g. in a *gdpP* mutant). Strikingly, the *agrA* mutation also significantly restored sensitivity to Congo Red to a *gdpP* mutant. We conclude that one or more genes under the direct or indirect control of AgrA block biofilm formation.

An important target of the *agr* operon is the effector molecule RNAIII. Indeed, the *gdpP* mutation markedly stimulated RNAIII transcript levels (Table 2). However, the known targets of RNAIII do not provide any obvious candidates for controlling cell lysis. The *agr* operon also controls transcription of a class of small proteins known as the phenol soluble modulins (PSMs). And indeed these modulins are reported to promote biofilm dispersal (Periasamy et al. 2012). However, their role in dispersal does not suggest a mechanism for how they could block biofilm formation and eDNA release. The PSM’s have also been reported to form amyloid fibers that stabilize *S. aureus* biofilms, but these fibers have not been detected in glucose-dependent biofilms (Schwartz et al. 2012). *In toto*, our results suggest that an as yet unidentified target(s) of *agr* is responsible for impeding biofilm formation until and unless c-di-AMP levels decrease.

An important unanswered question is how *agr* expression and that of genes under its control is regulated by c-di-AMP. While we do not have the answer, an appealing possibility is that the cyclic dinucleotide directly interacts with AgrA or some other protein product of the *agr* operon. Given that transcription of the operon is subject to positive autoregulation via quorum sensing, the stimulation of AgrA activity by c-di-AMP would stimulate transcription of the operon as well as that of downstream target genes (Ji et al. 1997; 1995).

Also remaining unanswered is the question of how the drop in c-di-AMP levels is brought about by fermentation during growth on glucose-containing medium. We found no significant difference in transcript levels of *dacA* or *gdpP* in response to glucose, suggesting that regulation of c-di-AMP occurs at a posttranscriptional level (Table S3). Previous studies have shown that the degradation product of c-di-AMP, 5’-phosphadenylyl-adenosine (pApA), as well as the alarmone (p)ppGpp are able to inhibit GdpP (Corrigan et al. 2015; Bowman et al. 2016), raising the possibility that the drop in c-di-AMP levels during biofilm formation is mediated at the level of the activity of GdpP.

Biofilm formation by *S. aureus* is critical to the formation of recalcitrant infections, such as infective endocarditis and postsurgical infections on indwelling medical devices (Bjarnsholt 2013; Wolcott et al. 2010; Moore 2017; Costerton 1999; Stewart and Costerton 2001). Once biofilms become established, they are resistant to antimicrobial treatments, making the infection difficult to eradicate. This antimicrobial resistance is complicated by the emergence of antibiotic resistant strains of *S. aureus* (MRSA and VRSA) (Koch et al. 2014). The identification of c-di-AMP phosphodiesterase as playing a critical role in biofilm formation raises the possibility of using it as a target for the discovery of drugs that block *S. aureus* biofilm formation. Importantly, many phosphodiesterase inhibitors have been developed to treat multiple conditions in humans (Abusnina and Lugnier 2017; Maehara et al. 2017; Zhou et al. 2017). Although these phosphodiesterases are structurally distinct from GdpP, the ability to create safe and effective drugs that target these enzymes brings merit to the idea of obtaining an inhibitor of a bacterial phosphodiesterase.

## Methods

### Strains and Growth Conditions

The bacterial strains used in this study are listed in Table S1. *S. aureus* were maintained in Tryptic Soy Broth (TSB) with no glucose [17 g/L of Peptone for Casein (BD); 3 g/L of Peptone from Soymeal (Amresco); 5 g/L of NaCl (Sigma); 2.5 g/L of dipotassium hydrogen phosphate (Macron)] or TSB with 0.5% glucose (TSBg). Unless noted, all experiments were performed in TSBg. Selection for drug resistance was maintained using 10 μg/mL Chloramphenicol for *S. aureus. gdpP*::TnΩ1 strains were made by transduction via ϕ85. Plasmids were incorporated using electroporation. Riboswich (pCDAribo) was synthesized by Integrated DNA Technologies (IDT; Coralville, Iowa) and moved into pASD132 using restriction cloning. The sequence of the riboswitch is (Kellenberger et al. 2015): AGCTTAGATCTCGATCCCGCGAAATTAATACGACTCACTATAGGGGCCCGGATAGC TCAGTCGGTAGAGCAGCGGCCGGATGTAACTGAATGAAATGGTGAAGGACGGGTCCAGCTTAATCAAACACGAACGGGGGAACCAACGATTGGCTGTTTTATTAACAGCCTTGGGGTGAATCTTACTAAGTAAGAGGGGGTACTCTGAATCCCTAATCCGACAGCTAACCTCGTAGGCTTGTTGAGTAGAGTGTGAGCTCCGTAACTAGTTACATCCGGCCGCGGGTCCAGGGTTCAAGTCCCTGTTCGGGCGCCATAGCATAACCCCTTGGGGCCTCTAAACGGGTCTTGAGGGGTTTTTTGCTCGAG

### Biofilm Assay

*S. aureus* overnight cultures were diluted 1:200 into 200 μl of fresh TSB or TSBg per well of a Nunc MicroWell 96-well plate (no. 167008; ThermoFisher) and incubated statically at 37°C for 24 hours. When specified 10 U of DNase I (04716728001; Roche) were added to the biofilms after 23 hours and were then returned to 37°C for 1 additional hour. The medium was then carefully removed and the biofilms were washed three times with 200 μl of PBS at pH 7.5. 100 μl of 0.1% Crystal Violet stain was then added to each well for 5 minutes. The Crystal Violet was then carefully removed and the biofilms were gently washed three times with 200 μl of PBS at pH 7.5. The stained biofilms were then dried at room temperature. Once dry, 200 μl of acidified ethanol was added to each well. 50 μl of the resuspended stained cells was then transferred to a fresh microtiter plate with 150 μl of PBS at pH 7.5 and the OD600 was immediately measured using a plate reader (Infinite 200 Pro, Tecan). The background value of stained wells without any inoculum was subtracted from the readings and averages and standard deviations were calculated. Biofilm biomass was calculated to be relative to the WT HG003 biofilm.

### eDNA Measurements

Biofilms were grown as described above. Once the biofilms were grown for 24 hours they were washed three times with 200 μl of PBS at pH 7.5 and then resuspended in 200 μl of PBS at pH 7.5 and transferred to a filter plate (0.2-μm AcroPrep Advance 96-well filter plates no. 8019; Pall). 100 μl of the filtrate was then combined with 100 μl of 2 μM SYTOX Green (no. S7020; ThermoFisher) in PBS. Fluorescence was then measured using a plate reader (Infinite 200 Pro, Tecan) with excitation and emission wavelengths of 465 nm and 510 nm respectively. The values from uninoculated wells were subtracted from the readings and averages and standard deviations were calculated. The amount of eDNA present was calculated to be relative to that of the WT HG003 biofilm.

### Visualizing Cell Clumping

Biofilms were grown as describes as above. After 24 hours of growth, cells were gently resuspended using one gentle aspiration and dispensation of the medium they were grown in. 1 μl of each sample was then placed on an agar pad and visualized using a Nikon Eclipse Ti inverted microscope equipped with an Orca R2 (Hamamatsu) camera and a 10×/0.30 Plan Fluor air objective (Nikon).

### Riboswitch assay

*S. aureus* strains containing pASD132 (empty vector) or pASD containing a c-di-AMP riboswitch (pCDAribo) were grown in TSB or TSBg with 20 μg/ml of chloramphenicol, 0.01% Trypan Blue (Invitrogen), and 250 μM of DFHBI-1T (No. 5610; Torcis) overnight shaking at 37°C in a plate reader (Tecan Infinite 200 Pro or Bio Tek Cytation5) with excitation and emission wavelengths of 460 nm and 510 nm, respectively, for 16 hours. At each time point the empty vector fluorescence was subtracted from the pCDAribo strains to remove autofluorescence. Excluding Figure 3A the baseline for the assays was normalized to HG003 grown with no added glucose. Values are reported as arbitrary units (AU).

### Congo Red Assay

10-fold serial dilutions of overnight cultures were plated onto TSB, TSB + 0.1% Congo Red and 0.25% Congo Red. Plates were incubated overnight at 37°C prior to imaging and contained 20 μg/ml of chloramphenicol to maintain plasmids if needed.

### qPCR

*S. aureus* overnight cultures were diluted 1:200 into 3ml of fresh TSB or TSBg per well of a Nunc MicroWell 6-well plate (no. 140675; ThermoFisher) and incubated statically at 37°C for 6 hours. 6ml of RNA Protect Bacterial Reagent was added to each sample and biofilms were immediately resuspended by pipetting. The cells were pelleted at 5k x g for 5 minutes and resuspended in 1ml Trizol. Cells were then mechanically disrupted using a FastPrep-24 (MP Biomedicals). Chloroform extraction of trizol was done to isolate RNA. RNA was incubated with 10U DNase I (Roche) for 90 minutes at 37C to remove any DNA contamination followed by phenol/chloroform extraction to purify the RNA. cDNA was made using the Superscript IV first-strand synthesis system (18091050, ThermoFisher). qPCR was performed using QuantiTect SYBR Green (204145, Qiagen) and run on a BioRad CFX384. Data was analyzed using BioRad CFX analyzer 3.1. Data was normalized to transcript levels of the reference gene SAOUHSC_01490 (*hu*). Primers were designed using Geneious 11.0.5. Primers for qPCR are listed in Table S6.

## Acknowledgments

We thank former members of the Losick laboratory N. Bradshaw, A. DeFrancesco, and V. Haunreiter for discussions. We also thank A. Gründling for advice on the manuscript. We also thank S. Trauger and C. Vidoudez at the Harvard Small Molecule Mass Spectrometry core for the HPLC MS/MS work. This work was supported by the NIH grant AI139011 to RL and SW and GM18568 to RL.

