## Supplemental tables and figures for "Biofilm Formation by *Staphylococcus aureus* is Triggered by a Drop in the Levels of the Second Messenger cyclic-di-AMP"

Table S1. Bacterial strains used in this study

| **Name** | **Species** | **Strain background** | **Genotype/plasmid** | **Source** |
| --- | --- | --- | --- | --- |
| AS0065 | *S. aureus* | HG003 |  | (Herbert et al. 2010) |
| AS0069 | *S. aureus* | SH1000 |  | (O'Neill 2010) |
| AS0070 | *S. aureus* | LAC |  | (Voyich et al. 2005) |
| AS0192 | *S. aureus* | Newman |  | (Duthie and Lorenz 1952) |
| AS0194 | *S. aureus* | MN8 |  | (Maira-Litrán et al. 2002) |
| AS0195 | *S. aureus* | RN1 |  | (Herbert et al. 2010) |
| AS0133 | *S. aureus* | HG003 | *gdpP*::TnΩ1 | (DeFrancesco et al. 2017) |
| AS0218 | *S. aureus* | SH1000 | *gdpP*::TnΩ1 | This work |
| AS0219 | *S. aureus* | LAC | *gdpP*::TnΩ1 | This work |
| AS0220 | *S. aureus* | Newman | *gdpP*::TnΩ1 | This work |
| AS0222 | *S. aureus* | MN8 | *gdpP*::TnΩ1 | This work |
| AS0223 | *S. aureus* | RN1 | *gdpP*::TnΩ1 | This work |
| AS0144 | *S. aureus* | HG003 | pASD132 (empty vector) | (DeFrancesco et al. 2017) |
| AS0145 | *S. aureus* | HG003 | pCDAribo | This work |
| AS0244 | *S. aureus* | HG003 | pASD133 (*gdpP)* | (DeFrancesco et al. 2017) |
| AS0150 | *S. aureus* | HG003 | *gdpP*::TnΩ1 + pASD132 | (DeFrancesco et al. 2017) |
| AS0151 | *S. aureus* | HG003 | *gdpP*::TnΩ1 + pCDAribo | This work |
| AS0211 | *S. aureus* | HG003 | *gdpP*::TnΩ1 + pASD133 | (DeFrancesco et al. 2017) |
| AS0725 | *S. aureus* | HG003 | *gdpP*::TnΩ1 + pASD133D418A | This work |
| AS0152 | *S. aureus* | HG003 | ∆*xdrA* + pASD132 | This work |
| AS0153 | *S. aureus* | HG003 | ∆*xdrA* + pCDAribo | This work |
| AS0285 | *S. aureus* | HG003 | ∆*xdrA* + pASD133 | This work |
| AS0726 | *S. aureus* | HG003 | ∆*xdrA* + pASD133 D418A | This work |
| AS0154 | *S. aureus* | HG003 | ∆*apt* + pASD132 | This work |
| AS0155 | *S. aureus* | HG003 | ∆*apt* + pCDAribo | This work |
| AS0286 | *S. aureus* | HG003 | ∆*apt* + pASD133 | This work |
| AS0416 | *S. aureus* | HG003 | ∆*xdrA/∆apt* + pASD132 | This work |
| AS0417 | *S. aureus* | HG003 | ∆xdrA/∆apt + pCDAribo | This work |
| AS0294 | *S. aureus* | HG003 | ∆xdrA/∆apt + pASD133 | This work |
| AS0212 | *S. aureus* | HG003 | Δ*agr* | This work |
| AS0213 | *S. aureus* | HG003 | *gdpP*::TnΩ1 /Δ*agr* | This work |
| AS0214 | *S. aureus* | HG003 | Δ*agr* + pASD132 | This work |
| AS0215 | *S. aureus* | HG003 | Δ*agr* + pCDAribo | This work |
| AS0216 | *S. aureus* | HG003 | *gdpP*::TnΩ1 /Δ*agr* + pASD132 | This work |
| AS0217 | *S. aureus* | HG003 | *gdpP*::TnΩ1 /Δ*agr* + pCDAribo | This work |

Table S2. HPLC MS quantification of c-di-AMP (pmol/OD)

| Plasmid | Glucose | HG003 | *gdpP*::TnΩ1 | ∆*xdrA* | ∆*apt* | ∆*xdrA*/∆*apt* |
| --- | --- | --- | --- | --- | --- | --- |
| pASD132 | - | 264 | 216 | 15 | 156 | 190 |
| pASD132 | + | 37 | 202 | 289 | 218 | 714 |
| pASD133 (*gdpP*) | + | 20 | 52 | 30 | 17 | 252 |

­­

pASD132 is the empty vector and pASD133 carries the *gdpP* gene (Table S1).

HPLC MS was carried out as follows:

20 μl overnight culture was used to inoculate 10 mL of TSB or TSBg grown shaking at 37°C for 6 hours. 25 mL of OD_600_ = 1 were then washed once with ice-cold water. Cell pellets were then immediately resuspended in 500 μl of methanol (MX0486-1; Sigma) and transferred to an RNase and DNase free bead beater tube containing Lysing Matrix B (116911500, MP Biomedicals). Collection tubes were washed with 500 μl of methanol and pooled with the rest of the sample. Samples were run twice on setting 6 for 1 minute on an FastPrep -24 Homogenizer (116004500, MP Biomedicals). Tubes were then centrifuged for 5000 x g for 30 seconds to pellet beads and cell debris. Supernatants were transferred to a glass vial (66011-085; VWR). Beads were then washed with 1 mL of methanol and supernatant transferred to the glass vial. 4 mL of ice-cold chloroform was added and each sample was vortexed for 1 minute. 2 mL water (WX0001-1; Sigma) which contained 10 pM/mL of c-di-GMP (SML1228-1UMO; Sigma) as an internal standard was then added and samples were vortexed for 1 minute. Vials were then centrifuged for 10 minutes at 3000 rpm. The aqueous top phase was transferred to a new glass vial, being careful not to transfer any of the interphase, and dried under N2. Once dry samples were stored at -80°C until HPLC. All samples were run on a ThermoFisher Q-exactive with a Restek PFPP 150x2 mm column maintained at 25°C. Samples were separated at a flow rate of 0.2 mL/min using the following buffers and gradient: buffer A (20 mM of ammonium acetate, 0.1% formic acid in water) and buffer B (methanol, 0.1% formic acid) for 0–4 min (0% buffer B), 4–9 min (100% buffer B), 9–11 min (0% buffer B). The MS parameters are as follows: Polarity +, Full MS, Resolution 70000, AGC target 3e6, mz range: 300 to 700; Polarity - Full MS, Resolution 70000, AGC target 3e6, mz range: 300 to 700. A standard curve of c-di-AMP was made using c-di-AMP sodium salt (SML1231-1UMO; Sigma).

Table S3. Transcript levels of *dacA* and *gdpP* under biofilm forming conditions (presence of glucose)

| **Locus Tag** | **Gene** | **Fold change** |
| --- | --- | --- |
| SAOUHSC_02407 | *dacA* | -0.3 ± 0.8 |
| SAOUHSC_00015 | *gdpP* | 2.2 ± 0.4 |

Table S4: Transcript levels of *dacA* and *gdpP* under biofilm forming conditions compared to HG003

| **Strain** | ***dacA* (SAOUHSC_02407)** | ***gdpP* (SAOUHSC_00015)** |
| --- | --- | --- |
| *gdpP*::TnΩ1 | 2.0 ± 0.3 | - |
| ∆*xdrA* | -0.5 ± 0.9 | 0.3 ± 0.8 |
| ∆*apt* | -1.8 ± 0.4 | 3.7 ± 0.8 |
| ∆*xdrA*/∆*apt* | -0.1 ± 1.4 | 3.1 ± 1.4 |

qPCR Method: qPCR as described in main text. Samples were run in technical triplicate for three biological replicates. Triplicate Cq data was averaged to use in ΔΔCt analysis. The table of results is the average of the 3 biological replicates for that sample type. Error is calculated as the standard error from the biological replicate averages.

Table S5. Transposon insertions in genes that lead to Congo Red Resistance

| **Functional category** | **Locus tag** | **Gene** | **Product** | **Ratio** | **Underrepresented in eDNA** |
| --- | --- | --- | --- | --- | --- |
| Cell wall-related | SAOUHSC_02885 | *oatA* | O-acetyltransferase | 12.8206 |  |
| Cell wall-related | SAOUHSC_02012 | *sgtB* | Glycosyltransferase | 10.1897 |  |
| Cell wall-related | SAOUHSC_01359 | *fmtC* | Phosphatidylglycerol lysyltransferase | 6.1699 |  |
| Electron transport chain | SAOUHSC_00982 | *menF* | Menaquinone biosynthesis | 25.3775 |  |
| Electron transport chain | SAOUHSC_01618 | *ispA* | Geranyltranstransferase | 23.3373 |  |
| Electron transport chain | SAOUHSC_01487 | *ubiE* | Menaquinone biosynthesis | 20.3097 |  |
| Electron transport chain | SAOUHSC_01916 | *menE* | Menaquinone biosynthesis | 15.9142 |  |
| Electron transport chain | SAOUHSC_01482 | *aroB* | 3-dehydroquinate synthase | 14.2334 |  |
| Electron transport chain | SAOUHSC_00983 | *menD* | Menaquinone biosynthesis | 12.1464 |  |
| Electron transport chain | SAOUHSC_01483 | *aroC* | Chorismate synthase | 8.9793 |  |
| Electron transport chain | SAOUHSC_01481 | *aroA* | 3-phosphoshikimate 1-carboxyvinyltransferase | 6.7209 |  |
| Electron transport chain | SAOUHSC_01852 | *aroA2* | Chorismate mutase | 6.6593 |  |
| Electron transport chain | SAOUHSC_00832 | *aroD* | 3-dehydroquinase | 5.3866 |  |
| Electron transport chain | SAOUHSC_01960 | *hemY* | Protoporphyrinogen oxidase | 19.481 |  |
| Electron transport chain | SAOUHSC_01776 | *hemA* | Glutamyl-tRNA reductase | 11.6706 |  |
| Electron transport chain | SAOUHSC_01961 | *hemH* | Ferrochelatase | 8.3048 |  |
| Electron transport chain | SAOUHSC_01962 | *hemE* | Uroporphyrinogen decarboxylase | 6.2842 |  |
| Electron transport chain | SAOUHSC_01065 | *ctaA* | Heme A synthase | 5.1375 |  |
| Glycolysis/TCA cycle | SAOUHSC_00798 | *pgm* | Phosphoglycerate mutase | 15.3775 |  |
| Glycolysis/TCA cycle | SAOUHSC_02366 | *fbaA* | Fructose-bisphosphate aldolase | 8.3939 |  |
| Nucleotide metabolism | SAOUHSC_00372 | *xpt* | Xanthine phosphoribosyl transferase | 11.5975 |  |
| Nucleotide metabolism | SAOUHSC_01742 | *relA* | GTP pyrophosphokinase | 9.6613 |  |
| Nucleotide metabolism | SAOUHSC_01251 | *pnpA* | Polynucleotide phosphorylase | 8.8043 |  |
| Nucleotide metabolism | SAOUHSC_00015 | *gdpP* | c-di-AMP phosphodiesterase | 7.4069 | * |
| Nucleotide metabolism | SAOUHSC_00374 | *guaB* | IMP dehydrogenase | 6.7569 |  |
| Protease | SAOUHSC_01778 | *clpX* | ATP-dependent protease | 9.0337 |  |
| Transcription | SAOUHSC_02369 | *rpoE* | RNA polymerase subunit delta | 22.1268 |  |
| Transcriptional Regulator | SAOUHSC_01979 | *xdrA* | Transcriptional regulator | 60.2878 | * |
| Transcriptional Regulator | SAOUHSC_01586 | *srrA* | DNA-binding response regulator | 24.3428 | * |
| Transcriptional Regulator | SAOUHSC_02664 | *rsp* | Transcriptional regulator | 5.2234 |  |
| Transporter | SAOUHSC_00420 | *-* | Sodium-dependent transporter | 11.7191 | * |
| Transporter | SAOUHSC_00373 | *pbuX* | Xanthine permease | 10.6772 |  |
| Transporter | SAOUHSC_00636 | *mntB* | Manganese transporter | 6.5823 |  |
| Transporter | SAOUHSC_00637 | *mntA* | Manganese transporter | 6.4466 |  |
| Transporter | SAOUHSC_00634 | *mntC* | Manganese transporter | 6.2462 |  |
| Unknown | SAOUHSC_00755 | *-* |  | 31.9346 | * |
| Unknown | SAOUHSC_00938 | *-* |  | 19.2136 |  |
| Unknown | SAOUHSC_00965 | *-* |  | 13.0967 |  |
| Unknown | SAOUHSC_02896 | *-* |  | 11.7309 |  |
| Unknown | SAOUHSC_00014 | *yybS* |  | 11.6834 | * |
| Unknown | SAOUHSC_01265 | *-* |  | 5.736 |  |
| Unknown | SAOUHSC_01076 | *-* |  | 5.4735 |  |

The screen for insertions conferring Congo Red resistance was carried out as follows:

Transposon sequencing of HG003 treated with Congo Red was performed as previously described (Santiago et al. 2015). A pooled transposon library containing ~ 5 x 10^5^ cells was grown both in the presence and absence of 16 µg/mL Congo Red at 37°C until the OD600 reached 1.0. Genomic DNA was isolated and prepared for Illumina sequencing as previously described. Data was processed using the Galaxy webserver. Transposons were mapped to the *S. aureus* NCTC8325 genome using Bowtie (Langmead et al. 2009). The Mann–Whitney U test was used to identify genes with enriched reads over the control (Santiago et al. 2018). The threshold for significance was set to q < 0.05, where q is the corrected p-value. Genes that had more than a five-fold increase in the number of transposon insertions between the untreated control and Congo Red treated samples were considered to be significantly enriched.

Table S6. qPCR primers used in this study.

| **Target** | **Sequence** | **Source** |
| --- | --- | --- |
| SAOUHSC_01490 (*hu*) | TTTACGTGCAGCACGTTCAC | (DeFrancesco et al. 2017) |
| SAOUHSC_01490 (*hu*) | AAAAAGAAGCTGGTTCAGCAGTAG | (DeFrancesco et al. 2017) |
| SAOUHSC_02407 (*dacA*) | TCGTTTGAAATGTCTCGTCGT | This study |
| SAOUHSC_02407 (*dacA*) | ACAAGACATAGAGCTGCGGT | This study |
| SAOUHSC_00015 (*gdpP*) | GAAAATTCTAAACCAATCATTGCGACA | This study |
| SAOUHSC_00015 (*gdpP*) | GCCCATCGACTAATGACACG | This study |
| SAOUHSC_02265 (*agrA*) | TCCTTATGAGGTGCTTGAGCA | This study |
| SAOUHSC_02265 (*agrA*) | CTGGGTCATGCTTACGAATTTCA | This study |
| RNAIII | AAGCCATCCCAACTTAATAACC | (Canovas et al. 2016) |
| RNAIII | GCACTGAGTCCAAGGAAACTAAC | (Canovas et al. 2016) |
| *psmα* | CCATGTGAATGGCCCCCTTC | This study |
| *psmα* | TCACATGGGTATCATTGCAGG | This study |

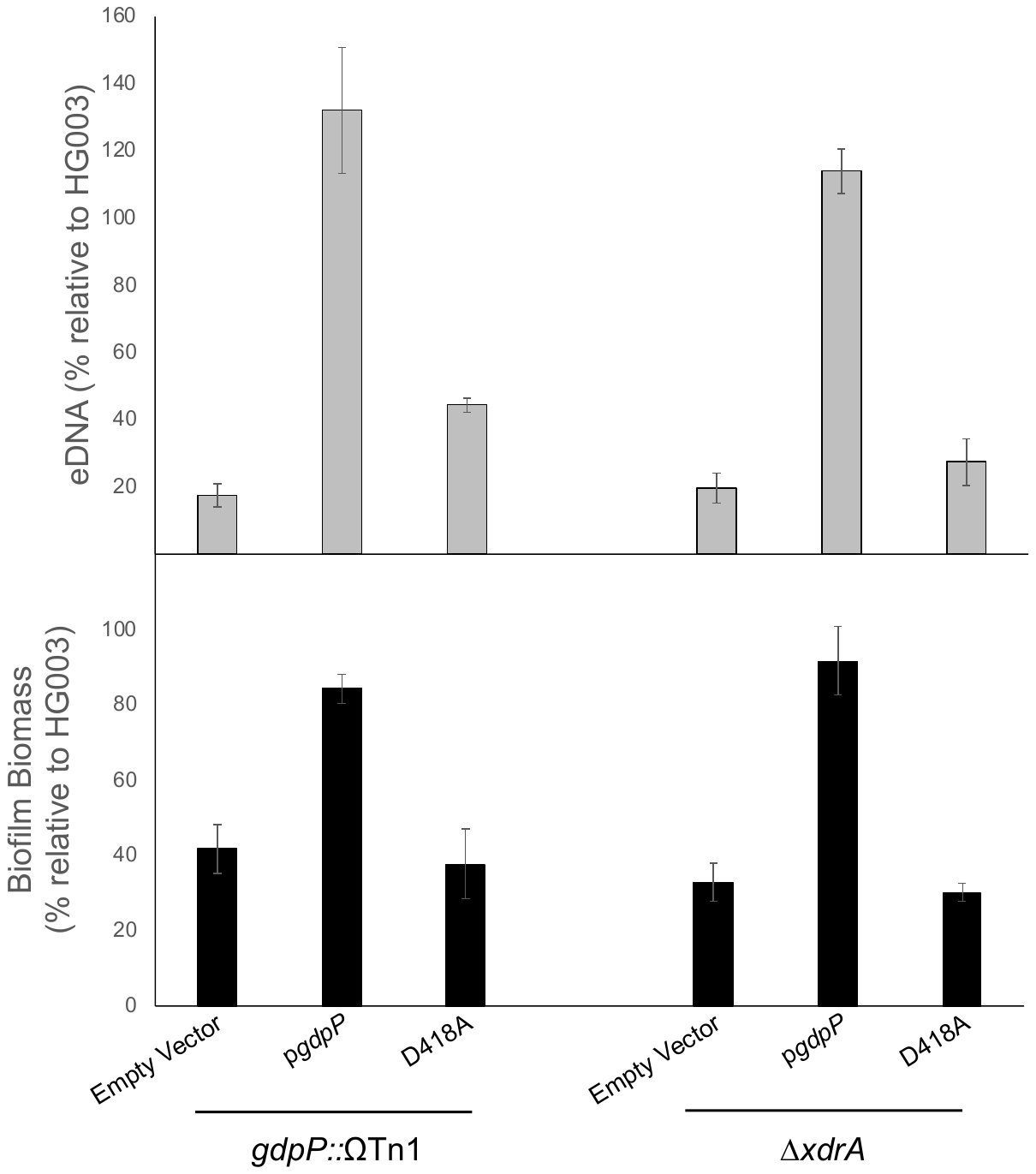

**Figure S1. Restoration of eDNA release and biofilm formation by overexpressing *gdpP* is dependent on its phosphodiesterase activity.** eDNA release (top) and biofilm formation (bottom) of the parental strains containing the empty vector plasmid (pASD132), a plasmid overexpressing *gdpP* (pASD133), or a plasmid overexpressing *gdpP* with a D418A substitution (pASD133D418A) during glucose-dependent biofilm formation.
